# Experimentally-constrained biophysical models of tonic and burst firing modes in thalamocortical neurons

**DOI:** 10.1101/512269

**Authors:** Elisabetta Iavarone, Jane Yi, Ying Shi, Bas-Jan Zandt, Christian O’Reilly, Werner Van Geit, Christian A. Rössert, Henry Markram, Sean L. Hill

## Abstract

Somatosensory thalamocortical (TC) neurons from the ventrobasal (VB) thalamus are central components in the flow of sensory information between the periphery and the cerebral cortex, and participate in the dynamic regulation of thalamocortical states including wakefulness and sleep. This property is reflected at the cellular level by the ability to generate action potentials in two distinct firing modes, called tonic firing and low-threshold bursting. Although the general properties of TC neurons are known, we still lack a detailed characterization of their morphological and electrical properties in the VB thalamus. The aim of this study was to build biophysically-detailed models of VB TC neurons explicitly constrained with experimental data from rats. We recorded the electrical activity of VB neurons (N = 49) and reconstructed morphologies in 3D (N = 50) by applying standardized protocols. After identifying distinct electrical types, we used a multi-objective optimization to fit single neuron electrical models (e-models), which yielded multiple solutions consistent with the experimental data. The models were tested for generalization using electrical stimuli and neuron morphologies not used during fitting. A local sensitivity analysis revealed that the e-models are robust to small parameter changes and that all the parameters were constrained by one or more features. The e-models, when tested in combination with different morphologies, showed that the electrical behavior is substantially preserved when changing dendritic structure and that the e-models were not overfit to a specific morphology. The models and their analysis show that automatic parameter search can be applied to capture complex firing behavior, such as co-existence of tonic firing and low-threshold bursting over a wide range of parameter sets and in combination with different neuron morphologies.

**Author summary:** Thalamocortical neurons are one of the main components of the thalamocortical system, which are implicated in key functions including sensory transmission and the transition between brain states. These functions are reflected at the cellular level by the ability to generate action potentials in two distinct modes, called burst and tonic firing. Biophysically-detailed computational modeling of these cells can provide a tool to understand the role of these neurons within thalamocortical circuitry. We started by collecting single cell experimental data by applying standardized experimental procedures in brain slices of the rat. Prior work has demonstrated that biological constraints can be integrated using multi-objective optimization to build biologically realistic models of neuron. Here, we employ similar techniques as those previously employed, but extend them to capture the multiple firing modes of thalamic neurons. We compared the model results with additional experimental data test their generalization and quantitatively reject those that deviated significantly from the experimental variability. These models can be readily integrated in a data-driven pipeline to reconstruct and simulate circuit activity in the thalamocortical system.

## Introduction

Thalamocortical (TC) neurons are one of the main components of the thalamus and have been extensively studied *in vitro* and *in computo*, especially in first order thalamic nuclei in different species (1). One of these nuclei, namely the ventral posterolateral nucleus (VPL), relays somatosensory, proprioceptive, and nociceptive information from the whole body to the somatosensory (non-barrel) cortex (2). The VPL is located close to ventral posteromedial nucleus (VPM), which transmits information from the face to the barrel cortex. The VPL and VPM nuclei constitute the ventrobasal (VB) complex of the thalamus (3).

Despite its key role in sensory functions, a systematic characterization of the cellular properties of the VB complex is still missing. The morphologies of VPL neurons in adult rats were described in early anatomical studies but were limited to two-dimensional drawings of Golgi-impregnated cells (4). The general electrical properties of TC neurons maintained *in vitro* are known and similar in different thalamic nuclei and species with respect to the generation of two distinct firing modes, called tonic firing and low-threshold bursting (5–8). However, a systematic description on the electrical types in the VB thalamus in the rodents is still missing.

Collecting morphological and electrophysiological data, by following standardized experimental procedures, is essential for the definition of cells types and it is the first step to constraining computational models of single neurons (9,10). Although models of TC neurons have already been previously published, they typically were aimed at studying specific firing properties and their parameters were hand tuned to achieve the desired result (11–15).

The purpose of our study is to systematically define the morphological and electrical types by collecting *in vitro* experimental data and to constrain biophysically detailed models of VB TC neurons of the juvenile rat. To the best of our knowledge, automatic parameter search has not been applied, thus far, to capture complex firing behavior in thalamic neurons, in particular low-threshold bursting and tonic firing. We defined the electrical and morphological types of TC neurons through in vitro patch-clamp recordings and 3D morphological reconstructions. We then extended an existing method (16) to account for their distinctive firing properties. These electrical models (e-models) were constrained by the electrical features extracted from experimental data (9,17,18). Other experimental data were used to assess the generalization of the models to different stimuli and morphologies. We further performed a sensitivity analysis by varying each parameter at a time by a small amount and recording the resulting electrical features. This analysis provides an assessment of the robustness of the models and a verification that the selected features provide sufficient constraints for the parameters.

## Results

### Physiological and morphological characterization

We characterized TC neurons in slices of the rat VB thalamus, by combining whole-cell patch-clamp recordings, biocytin filling and 3D Neurolucida (MicroBrightField) reconstruction, along with anatomical localization in a reference atlas (19) (Fig 1).

**Figure 1:**
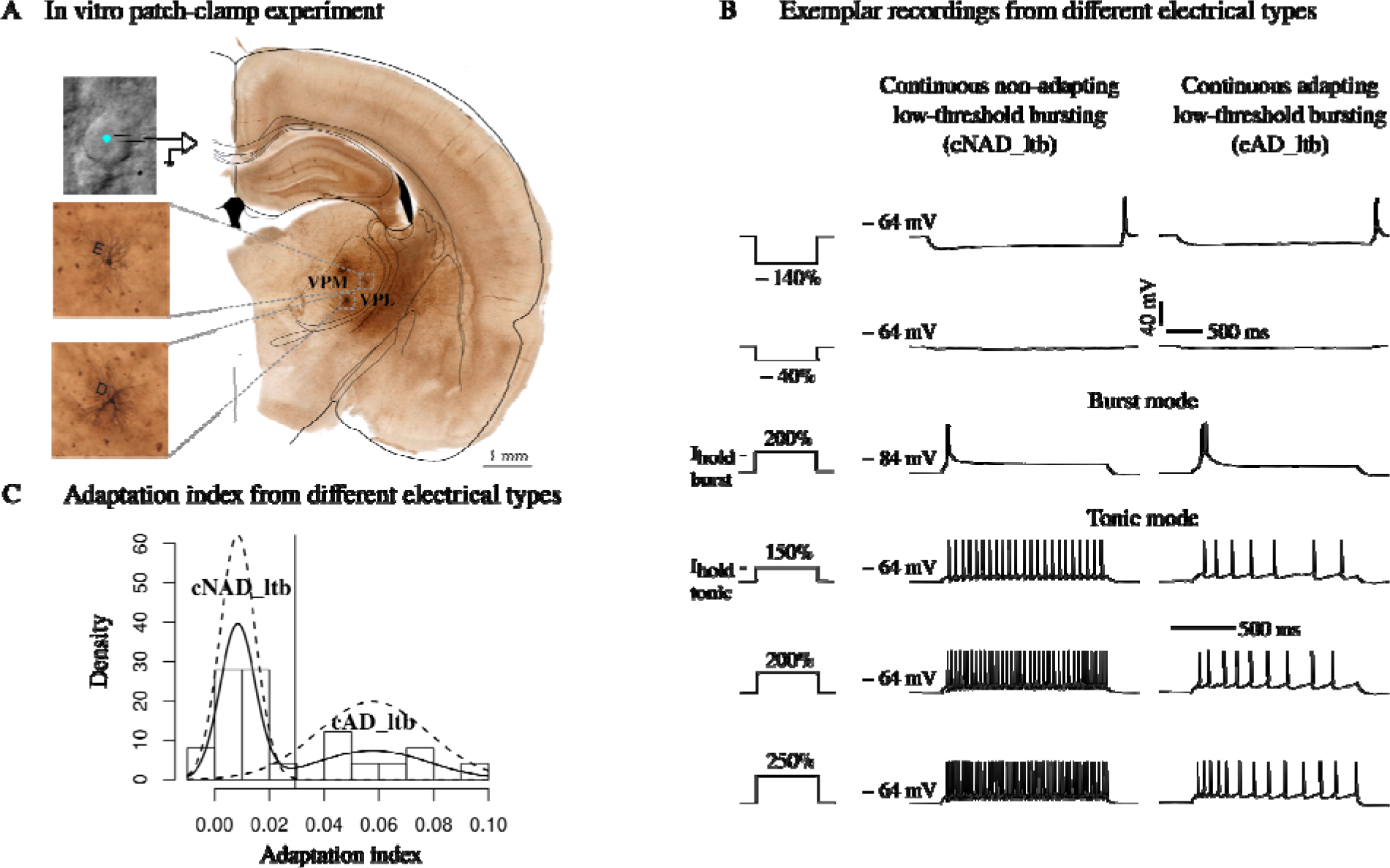
Simultaneous physiological and morphological characterization. (A) View of a patched cell under optic microscope and anatomical localization of biocytin-fille neurons (insets) in the rat Paxinos and Watson atlas (19). Letters D and E identify morphologie in a slice. (B) Voltage responses of two different thalamocortical (TC) neurons to a standardize battery of current stimuli. Each current amplitude was normalized by the threshold current of each neuron (e.g. 150 % threshold, see Methods). Third row is a low-threshold burst respons from a hyperpolarized holding potential, V_hold_ = −84 mV (burst mode), the other responses ar elicited from a depolarized holding potential, V_hold_ = −64 mV (tonic mode). Two differen holding currents (I_hold_ - tonic, I_hold_ - burst) are injected to obtain the desired V_hold_. The vertical scale bar applies to all the traces, the first horizontal scale bar from the top refers to the firs two rows, the second applies to the last four rows. (C) Analysis of adaptation index (AI) from recordings in tonic mode. Solid line is a non-parametric estimation of the distribution, dashed lines are two Gaussian distributions fitted to the data (see Methods). The vertical line indicate the cut-off value.

Visual inspection of 50 reconstructed morphologies (24 from the VPL, 26 from the VP nuclei) revealed variability in the number of principal dendritic trunks and their orientation, i agreement with previous anatomical studies (4).

The maximum radial extent of the dendrites ranged between 120 and 200 μm and they started to branch between 20 and 50 μm from the soma (Fig S1). We then analyzed the morphologies with two methods in order to quantitavely classify different morphological types. We used algebraic topology to extract the persistent homology of each morphology and to visualize the persistence barcode (20) (Fig 2A, see Methods). Each horizontal bar in the persistence barcode represents the start and end point of each dendritic component in terms of its radial distance from the soma. The barcodes of all the morphologies followed a semi-continuous distribution of decreasing length. To quantify the differences between the barcodes, we computed the pairwise distances of the persistence images (see Methods and Fig S1). We found that they were in general small (<0.4, values expected to vary between 0 and 1). These findings indicate that the morphologies cannot be grouped in different classes based on the topology of their dendrites. Furthermore, we performed Sholl Analysis (21) to compare the complexity of the dendritic trees (Fig 2B). We observed that all the morphologies had dense dendritic branches, with a maximum number of 50-100 intersections between 50-80 μm from the soma. When comparing the Sholl profiles for each pair of neurons we could not find any statistically significant difference (Fig S1C). Considering the results of topological and Sholl analyses, we grouped all the morphologies in one morphological type (m-type) called thalamocortical (TC) m-type.

**Figure 2:**
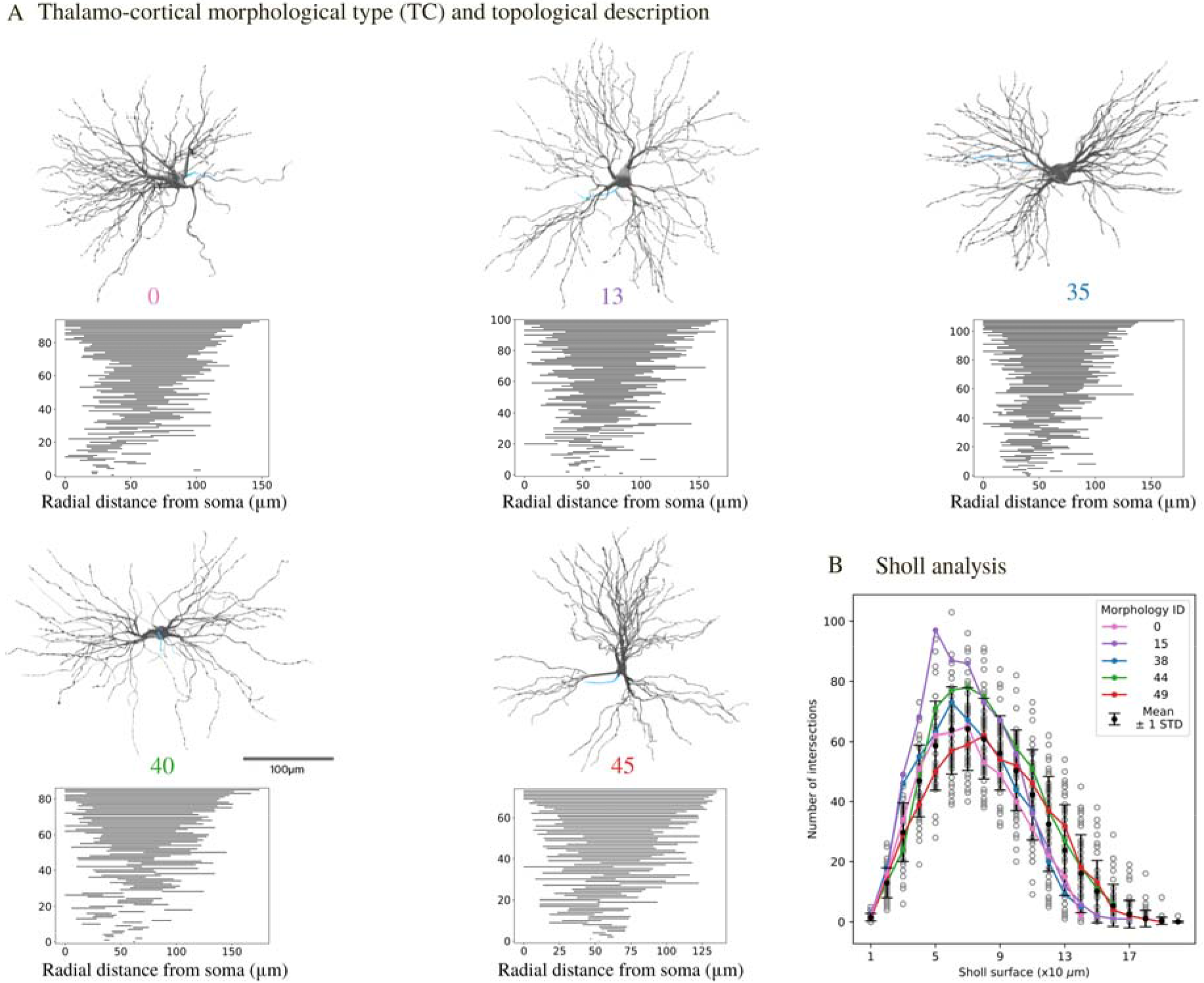
Morphological properties. (A) Renderings of 3D reconstructed TC neurons along with their persistence barcode according to (20). Grey: soma and dendrites, blue: axon only small sections available). The persistence barcode is a topological description of the branching pattern of the neurons’ dendrites. (B) Sholl analysis of TC neuron dendrites. For each Sholl ring, the number of intersections is shown (mean ± standard deviation, N = 50). Each grey circle represents one morphology, colored lines correspond to the morphologies in A. See Fig S1 for further analysis.

We used an adaptive stimulation protocol, called e-code, consisting of a battery of current stimuli (e- code, see Methods for details), where the stimulation amplitude was adapted to the excitability of different neurons. This standardized protocol has previously been used to build biophysically-accurate models of cortical electrical types (e-types) (16). However, TC neurons from different thalamic nuclei and species fire action potentials in two distinct firing modes, namely tonic firing, when stimulated from a relatively depolarized membrane potential or low-threshold bursting, from a hyperpolarized membrane potential (5). We thus extended the e-code to include two different holding currents. All the neurons recorded in this study displayed tonic and burst firing, when stimulated with the appropriate holding current (Fig 1). Moreover, we were able to classify different e-types by considering the voltage traces recorded in tonic mode in response to step current injections (Fig 1). The majority of the cells (59.3 %) showed a non-adapting tonic discharge (continuous non-adapting low-threshold bursting, cNAD_ltb e-type) while others (40.7 %) had higher adaptation rates (continuous non-adapting low-threshold bursting, cAD_ltb e-type), as reflected by the adaptation index (Fig 1C). We followed the Petilla convention (22) for naming the tonic firing discharge (cNAD or cAD), extending it to include “_ltb” for the low-threshold bursting property. In some rare examples, we noticed acceleration in the firing rate with decreasing inter-spike intervals (ISIs) towards the end of the stimulus. Similar adapting and accelerating responses have already been described in the VB thalamus of the cat (7). We also observed stereotypical burst firing responses within the same cell, with variation of the number of spikes per burst in different cells, but the burst firing responses alone were insufficient to classify distinct e-types.

### Constraining the models with experimental data

Multi-compartmental models comes with the need of tuning a large number of parameters (23), therefore we constrained the models as much as possible from experimental data. We first combined the morphology and the ionic currents models in the different morphological compartments (soma, dendrites and axon). Given that the reconstruction of the axon was limited, we replaced it with a stub representing the initial segment (16). We used previously published ionic current models and selected those that best matched properties measured in rat TC neurons (see Methods). The kinetics parameters were not part of the free parameters of the models. The distribution of the different ionic currents and their conductances in the dendrites of TC neurons is largely unknown. The current amplitudes of the fast sodium, persistent and transient (A-type) potassium currents were measured, but only up to 40-50 μm from the soma (24). Indirect measures of burst properties (15) or Ca^2+^ imaging studies (25) suggest that the low-threshold calcium (T-type) channels are uniformly distributed in the somatodendritic compartments. We thus assumed different peak conductance in the soma, dendrites and axon for all the ionic currents, except for *I*_*CaT*_, which had the same conductance value in the soma and dendrites. We then extracted the mean and standard deviation (STD) of different electrical features in order to capture the variability of firing responses from different cells of the same e-type (9) (Fig 3).

**Figure 3:**
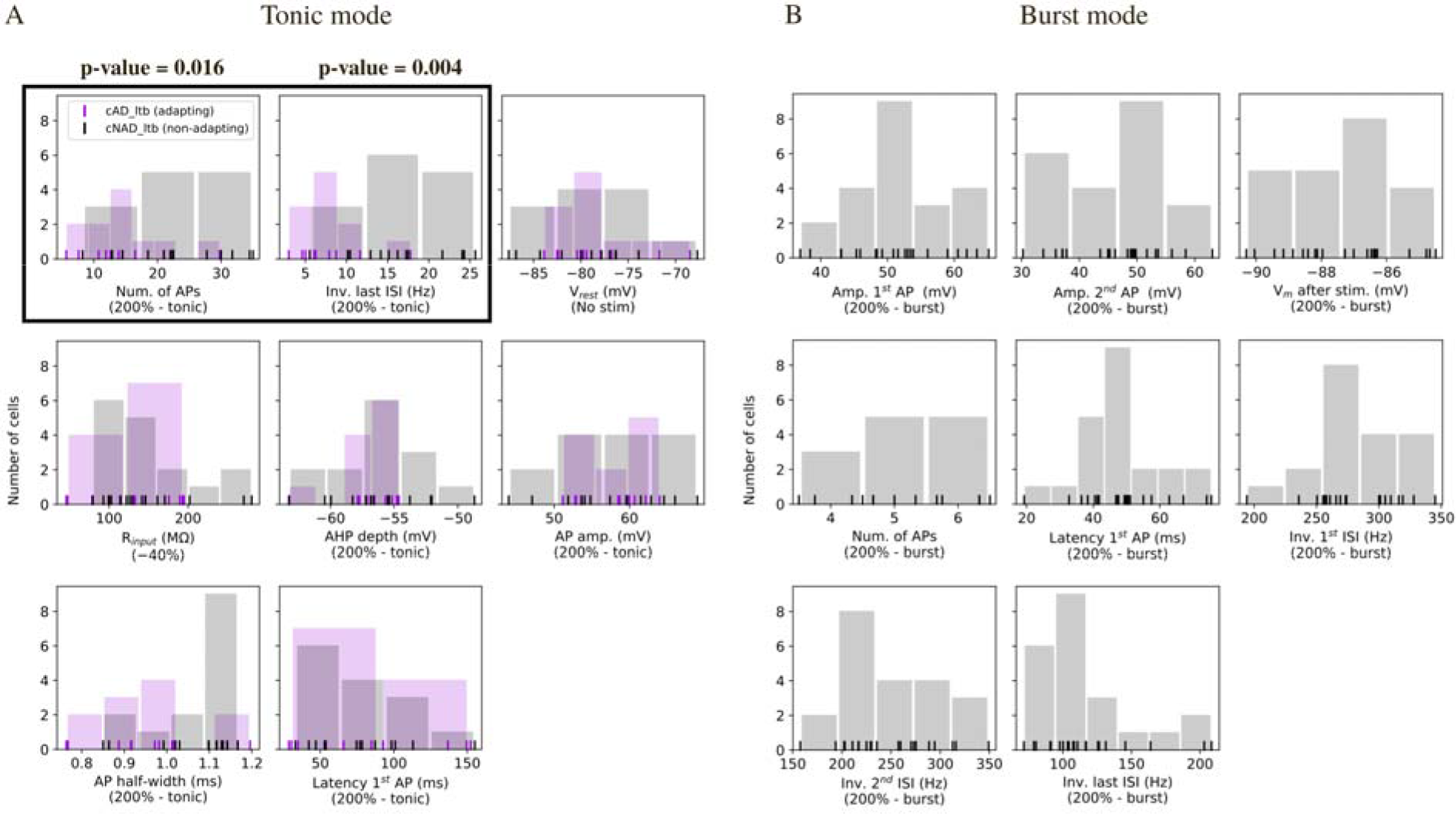
Histograms of electrical features. Each vertical line represents the mean feature value for a cell. Tonic and burst refer to th holding voltage as in Fig 1. (A) Feature values extracted from recordings in tonic mode (N = 1 cAD_ltb cells, N = 16 cNAD_ltb cells). The features highlighted by a black box show differen distributions for the cNAD_ltb and cAD_ltb electrical types (e-types) (p-value<0.05, two-side Mann-Whitney U test with Bonferroni correction for multiple comparisons). Passive propertie (V_rest_, R_input_) and spike shape features (AHP depth, AP amp., etc.) did not show clear difference between the two e-types. (B) Features measuring burst firing properties (N = 22 cells).

We observed that some features extracted from tonic firing responses had distinc distributions between the cAD_ltb and cNAD_ltb e-types (Fig 3A). The features were chosen i order to quantify salient physiological properties of TC neurons and to constrain the parameters of the model, namely the peak conductance of each ionic current. The average value and STD of the features were used as optimization objective (multi-objectiv optimization, MOO). Twenty-five parameters were allowed to vary between the upper an lower bounds shown in Fig 5. The models were associated with a training error, i.e. a set of all the feature errors (measured as absolute z-scores) (9,18,26)

**Figure 4:**
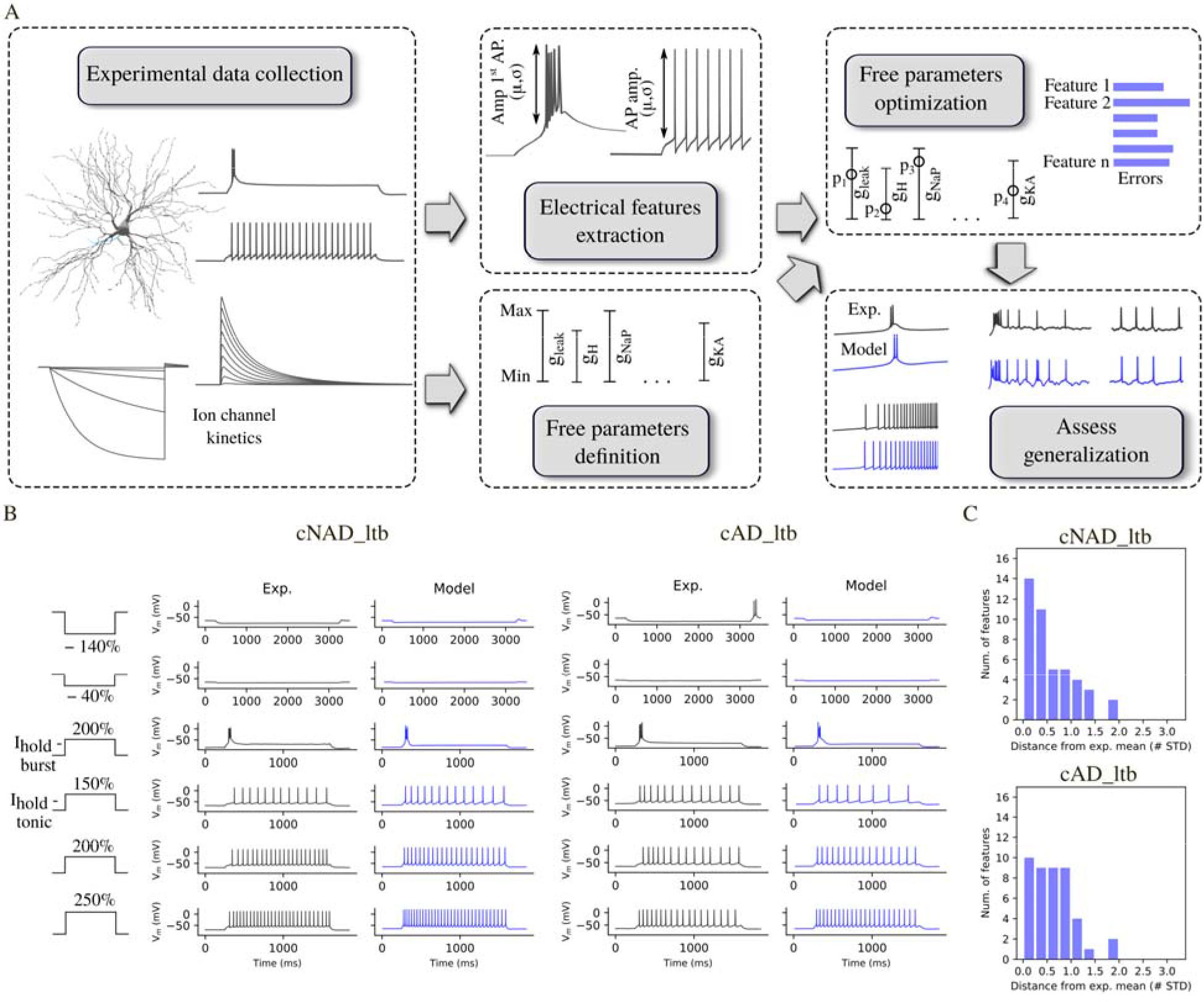
Models of different TC e-types and their fitting errors. (A) Single neuron modelling pipeline. (B) Experimental and model voltage responses to a variet of stimuli pattern used during the optimization of cNAD_ltb and cAD_ltb e-types. (C) Featur errors of the models shown in (B) reported as deviation from the experimental mean. The models are compared with the mean of features shown in Fig 3. Note that the models shown in B are fitted in order to reproduce the mean firing properties, not only a specific experimental recording. See Fig S2 for a complete list of fitting errors. By applying this MOO procedure, we generated multiple models with distinct parameter combinations that reproduced tonic and low-threshold burst firing in cNAD_ltb and cAD_ltb e-types (Fig 4).

### Model and experimental diversity

We found that different sets of parameter values reproduced the target firing behavior (Fig 5B). We further analyzed models that had all the feature errors below 3 STD. Models’ voltage responses reflected the characteristic firing properties of TC neurons (Fig S3), indicating that the selected set of features were sufficient to capture the two firing modes, in both the adapting and non-adapting e-types. The voltage traces from different models showed small differences in spike amplitude, firing frequency, and depth of the after-hyperpolarization, as reflected by the variability of features values (Fig 5C).

**Figure 5:**
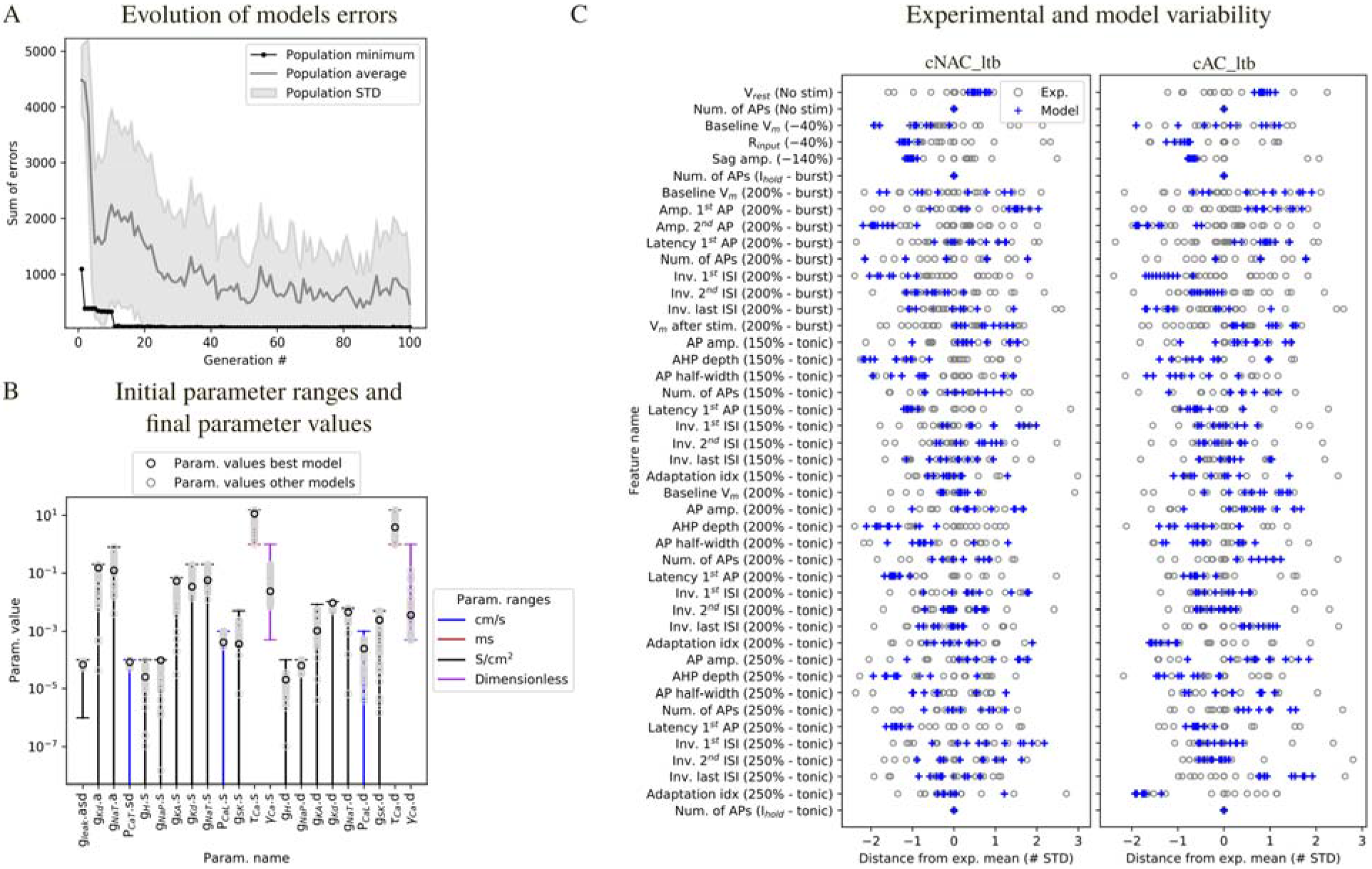
Diversity of model parameters and experimental variability. (A) Example of model fitting errors (sum of all feature errors) during optimization. (B) Initial parameter ranges and diversity of solutions. Each vertical line represents the range for th parameters, when the horizontal lower bar is missing the bound is 0. The characters followin “.” in the parameter name specifies the morphological compartment for the parameter (“s”: soma, “d”: dendrites, “a”: axon). Black circles: parameter values for one of the models in Fig 4 grey circles: parameter values of the models with all feature errors below 3 STD. (C) Feature variability in the models and experiments. Blue crosses: feature errors of a sample of 1 models. Each grey circle is the z-scored feature value of one experimental cell, obtained fro the feature values shown in Fig 3. The protocol names are shown in parenthesis an corresponds to the stimuli shown in Fig 1 and Fig 4, tonic and burst refer to the holding curren as in Fig 1.

Spike-shape related features (e.g. AP. amplitude) in the different models covered the space of the experimental variability, while for some features (e.g. input resistance, R_input_), all models tended to cluster on one of the tails of the experimental distribution. R_input_ relates to the neuron passive properties and depends both on the number of channels open at rest (inverse of the leak conductance in the model) and the size of the cell. Given that all the models were constrained on a single morphology, this result is not surprising. The number of action potentials (Num. of APs) in different conditions (No stim, I_hold_) ensured that the models did not spike in the absence of a stimulus or in response to the holding current. For this reason, all the experimental and model feature values in 5C are equal to 0. Other features, such as latency to the first spike and sag amplitude were less variable in the models compared to experiments. We hypothesized that this depended on the variable stimulation amplitudes applied to different experimental cells, while all the models were stimulated with the same current amplitudes.

We examined the diversity of the parameter values with respect to the initial parameter range (Fig 5B). Most of the optimized parameter values spanned intervals larger than one order of magnitude. On the other hand, some parameter values were restricted to one order of magnitude, for example the permeability of the low-threshold calcium current *P*_*CaT*_. This result is in agreement with experiments showing a minimum value of *I*_*CaT*_ is critical to generate burst activity and this critical value is reached only at a certain postnatal age (27). The value of *P*_*CaT*_ was constrained by features measuring burst activity (such as number of spikes, frequency, etc.).

### Assessment of model generalization

We used different stimuli for model fitting (current steps) and for generalizatio assessment (current ramps and noise). We simulated the experimental ramp currents in-silico, by stimulating the models with the appropriate holding currents for the two firing modes and linearly increasing current. We first compared visually the model responses with th experimental recordings (Fig 6A).

**Figure 6:**
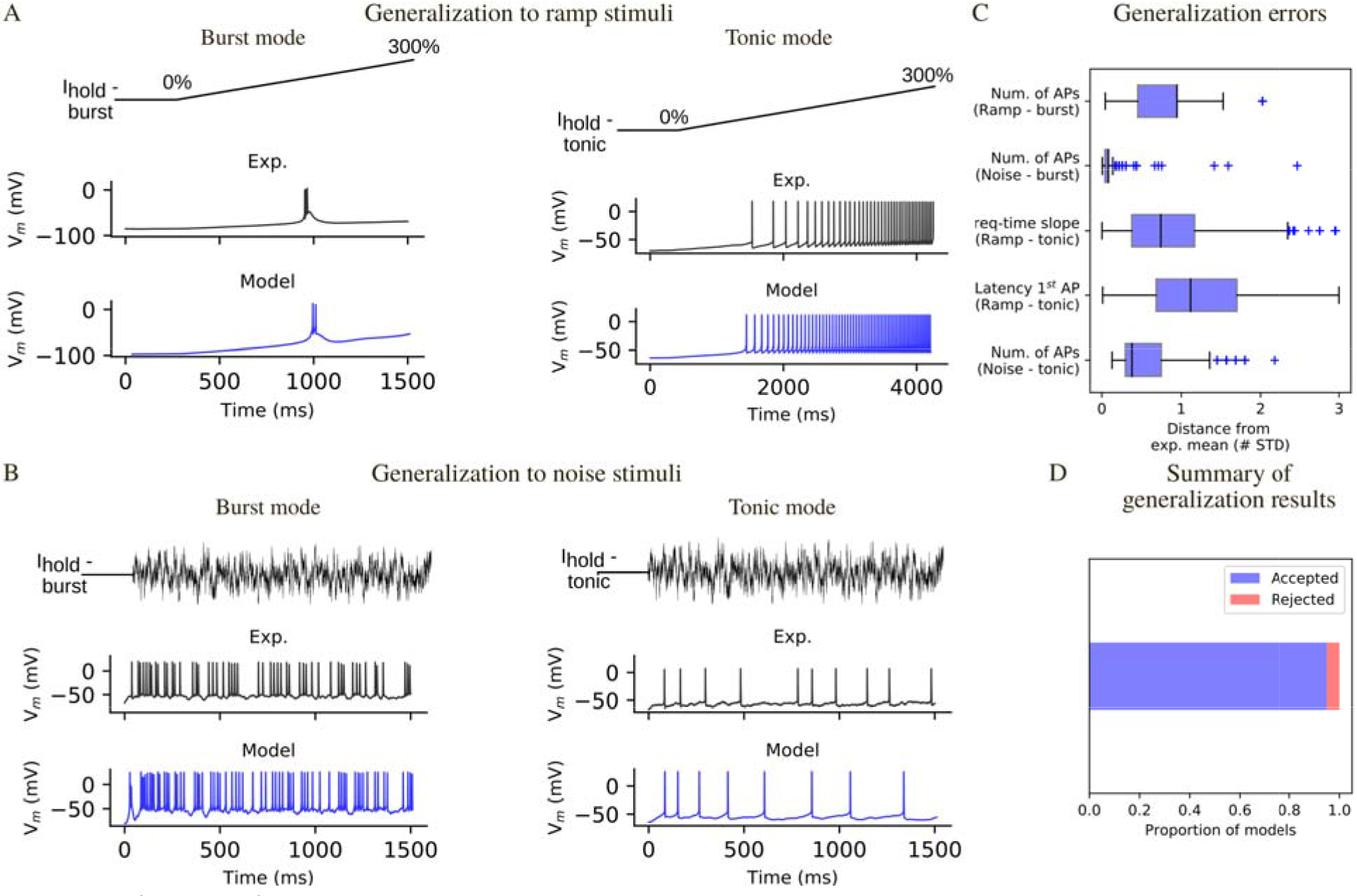
Model generalization. (A) Responses to a ramp current injection in burst mode (left) and tonic mode (center). (B Responses to a noise current generated according to an Ornstein-Uhlenbeck process and scale based on the excitability of the different experimental cells and models (see Methods). (C) Generalization errors for all the models that passed the generalization test (all generalization errors <3 STD). (D) Proportion of models that passed the generalization test (see Fig S4 for examples of models that failed this test).

In burst mode, the models reproduced the different behaviors observed experimentally: absence of a burst, small low-threshold spike, burst, burst followed by tonic firing (Fig S4). Moreover, the latency of burst generation substantially overlapped with the experimental one. However, a small fraction of models (1.2 %) generate repetitive burst that we have never observed in the experimental recordings (Fig S4). These models were quantitatively rejected by considering the number of spikes and the inter-spike intervals. In tonic mode, the latency to first spike, the voltage threshold, the shape of the subsequent action potentials and the increase in firing frequency were comparable with the experimental recordings (Fig 6A). In addition, we quantified the generalization error to ramp stimuli (Fig 6C), by considering the latency to first spike, firing frequency increase over time (tonic mode) or number of spikes (burst mode).

Although conductance-based models can be fit by using step and ramp currents (26), these stimuli are different from synaptic inputs, which can be simulated by injecting noisy currents. To test the response to such network-like input, we used a noisy current varying accordingly to an Ornstein-Uhlenbeck (OU) process (28) to compare models’ responses with the experimental data. Each experimentally recorded cell was stimulated with the same OU input, scaled by a factor *w*. Experimentally, *w* was calculated during the experiment by evaluating the responses to previous stimuli. We developed a similar approach to generate the noise stimuli in silico (see Methods). The noise current was injected on top of the holding currents used during the optimization. We found that the models reproduced well the subthreshold potential, spike times and the distribution of single spikes and bursts (Fig 6B). Moreover, we quantitatively evaluated the generalization to the noise stimulus by extracting features (e.g. number of spikes) and comparing them with the experimental mean.

We computed generalization errors for each model, which were calculated similarly to the optimization errors (Fig 6C). We considered a model acceptable after generalization if it had all generalization errors <3 STD and we found that the majority of the models (>90%) passed the generalization test.

### Sensitivity of electrical features to small parameter perturbations

We assessed the robustness of the models to small changes in their parameter values. To that end, we varied each parameter at a time by a small amount (± 2.5 % of the optimized value) and computed the values of the features. A sensitivity value of 2 between parameter p and feature y means that a 3 % change in p caused a 6 % change in f. We ranked the parameters from the most to the least influential and the features from the most sensitive to the least sensitive.

The conductance of the leak current *g*_*leak*_ emerged as the most influential parameter (Fig 7A). An increase in g_*leak*_ caused a decrease in firing frequency (inverse of inter-spike intervals, ISIs) i both the tonic and burst firing modes. These results are easy to interpret when considerin Ohm’s law: increasing *g*_*leak*_ means decreasing the input resistance of the model, so that for the same input current the voltage response becomes smaller. The second most influential parameter was the conductance of the persistent sodium current *g*_*NaP*_ in the dendrites, which increased the tonic firing rate as expected from a depolarizing current and had an effect on the late phase of the low-threshold burst (inverse last ISI - burst). An increase in the permeability of the low-threshold calcium current *P*_*CaT*_, known to be one the main currents underlying low threshold bursting, enhanced burst firing responses (it decreased the inverse of ISIs) and had effects on some of the tonic features. *P*_*CaT*_ was the third most influential parameter. These findings show that ICaT is the main driver of the low-threshold burst, but other currents, such as *I*_*NaP*_ contributes as well. Increasing the dendritic permeability of the high threshold calcium current *P*_*CaL*_ decreased the tonic firing rate, despite being a depolarizing current. Increasing P_*CaL*_ means higher Ca^2+^ influx and higher amplitude of the Ca^2+−^ activated potassium current (I_SK_). The parameter *g*_*SK*_ had indeed a similar effect on the features and thus clustered together with *P*_*CaL*_ (Fig 7B). Increasing the conductance of the transient sodium conductance *g*_*NaT*_ increased action potential amplitude and decreased its duration. Sag amplitude, that is known to depend on the activity of *I*_*H*_, was mainly influenced by change in *g*_*leak*_, *P*_*CaT*_ and *g*_*H*_. In summary, each parameter influenced at least one feature. Some features were weakly influenced by small parameter changes, e.g. baseline voltage, which depend more on the holding current amplitude, than on the model parameters. These results indicate that the model ability to generate tonic and burst firing is robust to small changes in parameter values and that all the parameters were constrained during the optimization by one or more features.

**Figure 7:**
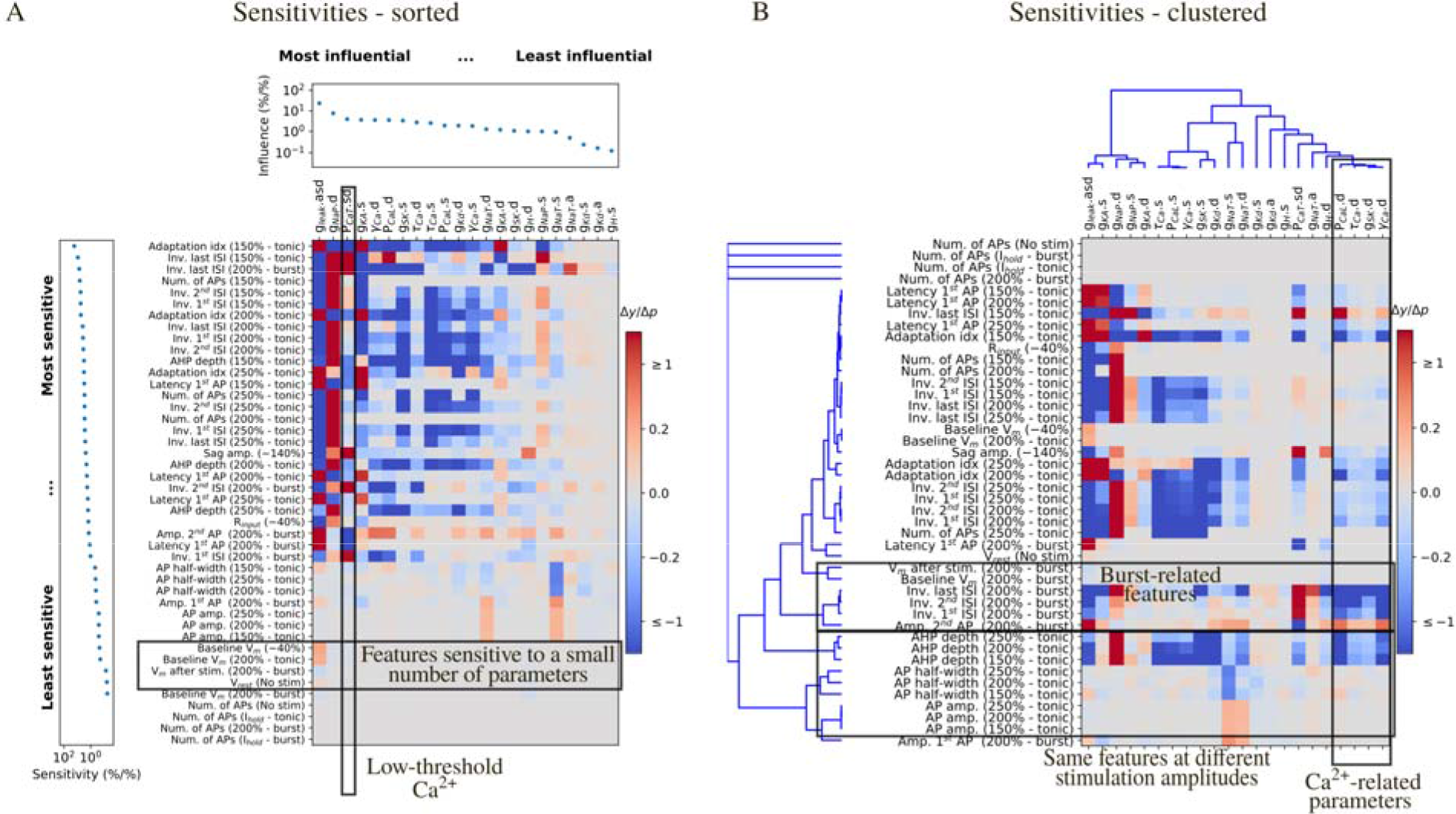
Local sensitivity analysis. (A) Sensitivity of the feature values to small changes to the parameter values for the cAD_lt model in Fig 4. Sensitivities (Δy/Δp) are color coded as a heat map. Features are ranked from the most to the least sensitive and parameters are ranked from the most to the leas influential. The last three rows are features that ensure that the models were not firing without input or in the response to the holding current. Small changes to the parameter values are not expected to make the model firing and thus the sensitivity of these features is 0. (B) Sam sensitivity values as in (a), with features and parameters clustered by similar sensitivity an influences.

We then analyzed which features depended similarly on parameter changes, as they may add superfluous degrees of freedom during parameters search. Fig 7B shows the same sensitivities as in Fig 7A, clustered by their similarities (see Methods). Features clustered together if they were sensitive to similar parameter combinations and parameters clustered based on their similar influence on the features. Not surprisingly, the same tonic features measured at different level of current stimulation clustered together (e.g. AP amplitude and half-width, AHP depth, latency of the first ISI) and tonic firing features belonged to a cluster that was different from burst features.

### Preservation of model firing properties with different morphologies

We optimized the parameters for the adapting and non-adapting e-models in combination with two different experimental morphologies and then tested them with the other 48 morphologies. Considering that morphologies could not be classified in different m-types based on topological analysis of their dendrites and that TC neurons have been shown to be electrically compact (15), we expected the electrical behavior to be conserved when changing morphology. Nonetheless, different neurons vary in their input resistance R_input_ and rheobase current Ithr due to variation in the surface area. Variation in R_input_ and I_thr_ made the current amplitude applied during the optimization inadequate to generate the appropriate voltage trajectories. We thus devised an algorithm to search for the holding current to obtain the target holding voltage (for example −64 mV or −84 mV for tonic and burst firing, respectively) and I_thr_ from the desired holding voltage. The different e-model/morphology combinations (me-combinations) were evaluated by computing the same feature errors calculated during optimization. For each morphology, we selected the e-model that generated the smalles maximum error. All me-combinations reproduced burst and tonic firing (Fig 8C). However, tw me-combinations generated responses with a small number of features that deviated from th experimental mean. We chose the value of 3 STD as a threshold to define which me combinations were acceptable (29), yielding 48 acceptable me-combinations out of the 5 tested (Fig 8A). We analyzed more closely which features were significantly different from th experimental mean. In Fig 8B we show that the rejected me-combinations had too many actio potentials in the burst.

**Figure 8:**
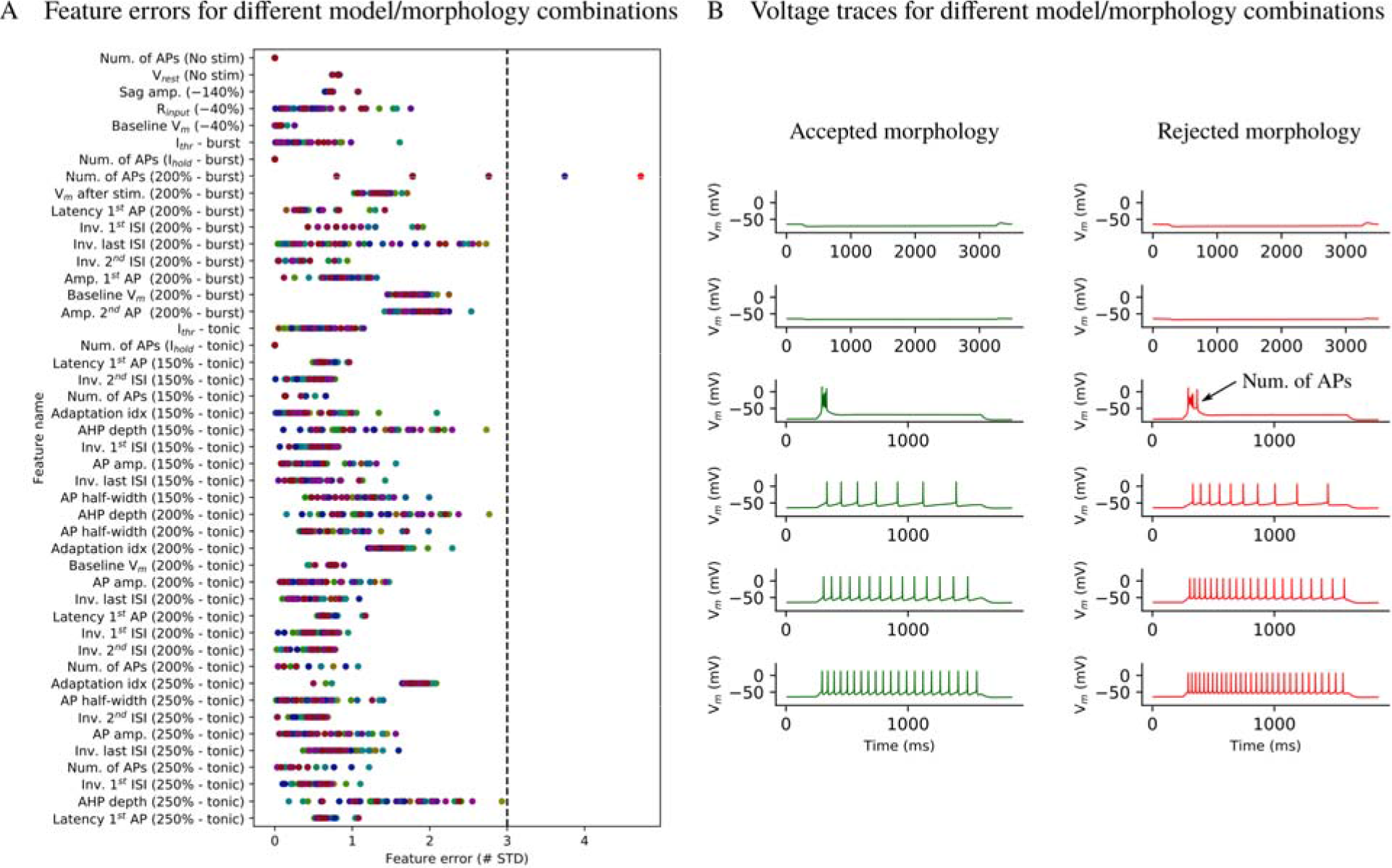
Model generalization to different experimental morphologies. (A) Feature errors from the best electrical models (e-model) showed in Fig 4 applied to 5 different TC cell morphologies. Each morphology is represented with a different color. E models/morphology combinations with at least one feature error > 3 STD (dashed line) were rejected. (B) Example of voltage responses from an accepted and from a rejected e- model/morphology combination. Feature errors for the rejected combination are shown in red in (A) and are indicated on the voltage trace.

## Discussion

Our objective was to apply and extend an existing data-driven pipeline to identify the cell types and build models of VB thalamocortical neurons that reproduce the multiple firing modes that have been experimentally observed. We successfully modelled these novel firing types, by including additional stimulation protocols and features to constrain the low-threshold burst.

Our morphological and electrical data were used to define the properties of VB TC neurons in the rat. We found two electrical types (e-types) of TC neurons, but no objectively different morphological types (m-types) were revealed either using Sholl analysis (21) or topological analysis of dendritic branching (20). We cannot exclude that refinements to these methods will reveal different m-types similar to the ones described in the visual thalamus of the mouse (30). We also showed that automatic parameter search can be applied to build biophysically and morphologically detailed models. This method was already applied to model canonical firing behavior in cortical, hippocampal and cerebellar granule neurons (9,10,16,17,31,32). To the best of our knowledge, such an automatic parameter search has not previously been used to capture different firing modes and complex firing behavior such as low-threshold bursting in thalamic neurons. Standardized electrophysiological protocols allowed us to identify for the first time in juvenile rat adapting and non-adapting e-types of TC VB neurons that were previously observed in other species (7). This finding suggests that the intrinsic properties of TC neurons contribut to adaptation, a key phenomenon for filtering out irrelevant stimuli, before sensory information reaches the neocortex. Further experiments are needed to elucidate the relative contribution of intrinsic mechanisms and network properties to adaptation in somatosensory systems. We named the two main e-types continuous non-adapting low-threshold bursting (cNAD_ltb) and continuous adapting low-threshold bursting (cAD_ltb) by following and extending existing conventions (16,22,31).

In this study, we improved upon previous morphologically and biophysically detailed models of tonic and burst firing in TC neurons (12,13,15) by explicitly constraining the parameters with experimental data, without hand-tuning of parameter values. Unlike previous models, we chose a multi-objective optimization for a methodological and a scientific reason: it is more time-efficient, reproducible, and it approximates the variability in ionic channel expression of biological neurons (31,33–35), as shown by the family of acceptable solutions we found. However, experiments aimed at quantifying ion channel conductances are essential to assess if these solutions fall between biological ranges. Furthermore, we tested the generalization capability of the models and found that more than 90% of the models were comparable with the experimental data.

Nonetheless, we noticed some inaccuracies when comparing the voltage traces with the experimental data when assessing the generalization of some models. For instance, some models tended to generate small transient oscillations in response to ramp stimuli in burst mode. This result is not surprising, considering that the exact kinetics for all the ionic currents are not available and that there are known limitations in models of ionic channels derived from the literature or from other models (36,37). In particular, modifications of the kinetics of the low-threshold calcium current was shown to explain the propensity to generate oscillatory bursts in TC neurons of other nuclei and species (38).

TC neurons have been shown to be electrically compact (15) and could, in principle, be modeled as a single compartment. However, active mechanisms need to be located in the dendrites in order to ensure synaptic integration and amplification (39). Information regarding specific conductances or firing properties in the dendrites of TC neurons is limited. For this reason, dendritic parameters in our models may be underconstrained. However, the sensitivity analysis (see below) revealed that dendritic parameters did not appear to be the least constrained because they influenced different tonic and burst-related features.

We included in the model fitting and validation pipeline a sensitivity analysis, which is often neglected in computational neuroscience (40). Although we cannot use our simple univariate approach to explore multidimensional parameter correlations and principles of co-regulation of ion channels expression, it is useful to find better constraints for parameters optimization. The selection of the features is indeed a step that still requires care and experience by modelers. Furthermore, this type of sensitivity analysis allows to identify parameters that can be traded-off during the optimization and that can be removed in order to reduce the dimensionality of the problem. In our study, four parameters related to the calcium dynamics were shown to influence the features in a very similar fashion. This type of analysis is of particular importance in future work aimed at using the full diversity of ion channels that can be inferred from gene expression data. More in detail, we propose that sensitivity analysis should be a fundamental tool in selecting which conductances are successfully optimized by the available experimental constraints. The example we showed is a local approach, applied to a specific solution to the optimization problem, which showed that our models are robust to small parameter changes. This analysis can be extended to study how the sensitivities vary in the neighborhood of different solutions.

In conclusion, we systematically studied the morphological and electrical properties of VB TC neurons and used these experimental data to constrain single neuron models, test their generalization capability and assess their robustness. Further work will validate these models in response to synaptic activity, in order to include them in a large-scale model of thalamocortical microcircuitry (16).

## Methods

### Experimental procedures

Experimental data were collected in conformity with the Swiss Welfare Act and the Swiss National Institutional Guidelines on Animal Experimentation for the ethical use of animals. The Swiss Cantonal Veterinary Office approved the project following an ethical review by the State Committee for Animal Experimentation.

All the experiments were conducted on coronal or horizontal brain slices (300 μm thickness) from the right hemisphere of male and female juvenile (P14-18) Wistar Han rats. The region of interest was identified using the Paxinos and Watson rat brain atlas (19). After decapitation, brains were quickly dissected and sliced (HR2 vibratome, Sigmann Elektronik, Germany) in ice-cold standard ACSF (in mM: NaCl 125.0, KCl 2.50, MgCl_2_ 1.00, NaH_2_PO_4_ 1.25, CaCl_2_ 2.00, D-(+)-Glucose 50.00, NaHCO_3_ 50.00; pH 7.40, aerated with 95% O_2_ / 5% CO_2_). Recordings of thalamocortical neurons in the VB complex were performed at 34 °C in standard ACSF with an Axon Instruments Axopatch 200B Amplifier (Molecular Devices, USA) using 5–7 MΩ borosilicate pipettes, containing (in mM): K^+^ -gluconate 110.00, KCl 10.00, ATP-Mg^2+^ 4.00, Na_2_-phosphocreatine 10.00, GTP-Na^+^ 0.30, HEPES 10.00, biocytin 13.00; pH adjusted to 7.20 with KOH, osmolarity 270-300 mOsm. Cells were visualized using infrared differential interference contrast video microscopy (VX55 camera, Till Photonics, Germany and BX51WI microscope, Olympus, Japan).

Membrane potentials were sampled at 10 kHz using an ITC-18 digitizing board (InstruTECH, USA) controlled by custom-written software operating within IGOR Pro (Wavemetrics, USA). Voltage signals were low-pass filtered (Bessel, 10 kHz) and corrected after acquisition for the liquid junction potential (LJP) of −14 mV. Only cells with a series resistance <25 MΩ were used.

After reaching the whole-cell configuration, a battery of current stimuli was injected into the cells and repeated 2-4 times (e-code). During the entire protocol, we defined offset currents in order to keep the cell at −50 mV (tonic firing) or −70 mV (burst firing) before LJP correction and applied them during the entire protocol. The step and ramp currents were injected with a delay of 250 ms in the experiment. In the models, the stimuli were injected with a delay of 800 ms, to allow for the decay of transients due to initialization. Each stimulus was normalized to the rheobase current of each cell, calculated on-line as the current that elicited one spike (stimulus TestAmp, duration 1350 ms). The stimuli used for in the experiments, for fitting and testing the models were:

- IDRest: current step of 1350 ms, injected at different amplitude levels in 25 % increments (range 50-300 % threshold). IDRest was renamed to Step in the model.
- IDThresh: current step with duration of 270 ms, 4 % increments (range 50 - 130 %).
- IV: hyperpolarizing and depolarizing steps of 3000 ms injected in 20 % increments (range 140 - 60%).
- SponNoHold: the first 10 seconds of this stimulus was used to calculate the resting membrane potential. No holding or stimulation currents were applied.
- SponHold: the first 10 seconds of this stimulus was used to calculate the holding current applied to keep the cells at the target potential.
- PosCheops: ramps of current from 0 to 300 % and from 300 to 0 % having progressively shorter durations (4000 ms, 2000 ms, 1250 ms). To test the models in tonic mode we used the first increasing ramp in the stimulus, while we used the last one in the bursting firing mode. We chose the last one because the biological cells were more likely to generate a burst.
- NOISEOU3: the original wave was scaled and offset for each cell based on the spike frequency responses to IDRest responses. The scaling factor w was extracted from the frequency-current curve and corresponded to the current value that made the cell fire at 7.5 Hz.

Neurons that were completely stained and those with high contrast were reconstructed in 3D and corrected for shrinkage as previously described (41). Reconstruction used the Neurolucida system (MicroBrightField). The location of the stained cells was defined by overlaying the stained slice and applying manually an affine transformation to the Paxinos and Watson’s rat atlas (19).

### Electrical features extraction

Electrical features were extracted using the Electrophys Feature Extraction Library (eFEL) (42). We calculated the adaptation index (AI) from recordings in tonic mode (Step 200 % threshold) and classified TC VB neurons into adapting (AI>=0.029) and non-adapting (AI<0.029) electrical types. AI was calculated using the eFEL *feature adaptation_index2* and corresponded to the average of the difference between two consecutive inter-spike intervals (ISI) normalized by their sum. The cut-off value was calculated after fitting a Gaussian mixture model to the bimodal data, using available routines for R (43,44). In order to group data from different cells and generate population features, we normalized all the stimuli by the rheobase current I_thr_ of each cell. To calculate I_thr_, we used IDRest and IDThresh and selected the minimal amplitude that evoked a single spike. The extracted features quantified passive (input resistance, resting membrane potential), burst and tonic firing properties (number of spikes, inverse of inter-spike intervals, latency to first spike), action potentials shape (amplitude, half-width, depth of the fast after-hyperpolarization). We aimed at finding the minimal set of features that capture the most important properties. This set was a trade-off between comprehensively describing the experimental data (i.e. extracting all possible features), which can lead to over-fitting and loss of generalizability, and a too small set that would miss some important characteristics. For the tonic firing responses, we used three stimulation amplitudes (150 %, 200 %, 250 % of firing threshold) which have been shown to reproduce the complete input-output function of the neurons (17,41). Responses to two hyperpolarizing steps of different amplitudes (−40 % and −140 % threshold) constrained the input resistance and the conductance of currents activated in hyperpolarization (*sag_amplitude* feature). We included baseline voltage values in the optimization objectives to ensure that the model was in the right firing regime and spike count to penalize models that were firing in response to the holding currents. Along with the voltage features, we extracted mean holding and threshold current values for all the experimental stimuli. Description of the features and the details on their calculation are available on-line [49]. Current stimuli applied during the optimization and generalization were directly obtained from the experimental values or automatically calculated by following the experimental procedures (e.g. noise stimulus).

### Morphology analysis

Reconstructed morphologies were analyzed to objectively identify different morphological types. The Sholl profiles of each pair of cells was statistically tested by using k-samples Anderson-Darling statistics. This test was preferred to the most common Kolmogorov-Smirnov test, because it does not assume that the samples are drawn from a continuous distribution. The different Sholl profiles are indeed an analysis of the intersections with discrete spheres.

To compare the topological description of each morphology we transformed the persistence barcodes into persistence images and calculated their distances as in (20). Briefly, we converted the persistence barcode, which encodes the start and end radial distances of a branch in the neuronal tree, into a persistence diagram. In the persistence diagram, each bar of the barcode is converted into a point in a 2D space, where the X and Y coordinates are the start and end radial distances of each bar. The persistence diagram was then converted in a persistence image by applying a Gaussian kernel. We used the library NeuroM (45) to perform Sholl and morphometrics analyses. The reconstructed morphologies will be made publicly available on neuromorpho.org.

### Ionic currents models

We used Hodgkin-Huxley types of ionic current models, starting from kinetics equations already available in the neuroscientific literature. Along with kinetics of the ionic currents, we stored information on the experimental conditions, such as temperature and LJP, by using the software NeuroCurator (46). Whenever the data was available, we compared simulated voltage-clamp experiments to experimental data from juvenile rats. Ionic currents *I*_*i*_ were defined as functions of the membrane potential *v*, its maximal conductance density gi and the constant value of the reversal potential *E*_*i*_:

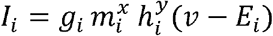

*m*_*ion*_ and *h*_*ion*_ represent activation and inactivation probability (varying between 0 and 1), with integer exponents *x* and *y*. Each probability varied according to:

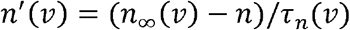

where *n*_*∞*_*(v)* is a function of voltage that represents the steady-state activation/inactivation function (normally fitted with a Boltzmann curve) and τ_*n*_*(v)* is a voltage-dependent time constant. Exceptions to this formalism are ionic currents that do not inactivate (y = 0) and ionic currents with (in)activation processes mediated by two or more time constants. Calcium currents (*I*_*CaT*_ and *I*_*CaL*_) were modeled according to the Goldman-Hodgkin-Katz constant field equation and had permeability values instead of conductance (47).

### Fast transient sodium current *I*_*NaT*_ and delayed potassium current *I*_*Kd*_

*I*_*NaT*_ and *I*_*Kd*_ were taken from a previous models of rat TC neurons from the VB nucleus (12), available on SenseLab ModelDB (accession no. 279). *I*_*NaT*_ was compared with recordings of transient sodium currents in P7-11 rat neurons from the dorsolateral geniculate (dLGN) nucleus (48).

### Low-threshold activated (T-type) calcium current *I*_*CaT*_

*I*_*CaT*_ model was taken from (12) and available on-line (ModelDB, accession no. 279). This model was based on data recorded from VB neurons of Sprague-Dawley rats (P7-12) at room temperature and corrected for −9 mV LJP (11).

### Hyperpolarization-activated cationic current *I*_*H*_

The steady-state activation for *I*_*H*_ was derived from VB thalamic neurons in P10-20 Long-Evans rats and was already corrected for −10 mV LJP in the original publication (49). The equation used was:

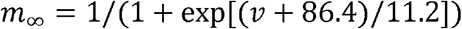

The time constant of activation was modeled as in (50), which derived a mathematical description of *I*_*H*_ based on data from the dLGN in adult guinea pigs, recorded at 35.5 °C (51). The equation describing the time dependence of activation was not corrected for simulations at different temperatures and was:

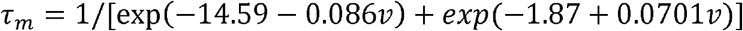

The equilibrium potential of the channel *E*_*H*_ was −43 mV. In silico voltage-clamp experiments were compared with data in (49).

### Persistent sodium current I_*NaP*_

We modeled I_*NaP*_ as in (17) which based their model on recordings from entorhinal neurons of Long-Evans rats (P25-P35) (52). The steady-state activation was modified according to (48) and the steady-state inactivation according to (14). The original steady-state activation data were recorded at room temperature (22-24°) and corrected for −6/−7 mV LJP. Limited data on *I*_*NaP*_ are available from dissociated neurons from the dLGN nucleus in Wistar rats (48).

### Fast transient (A-type) potassium current *I*_*KA*_

The mathematical formulation of I_KA_ was based on data recorded from VB neurons in Sprague-Dawley rats (P7-15), recorded at room temperature (22-24 °C) (53). A *Q*_*10*_ = 2.8 was experimentally determined and used for simulations at different temperatures. In the original experiments a small LJP (<−4 mV) was measured and not corrected. The current had a rapid and a slow component, represented by two activation and two inactivation variables. The model of this current was provided by the authors of (14).

### High-threshold (L-type) calcium current *I*_*CaL*_

*I*_*CaL*_ model is the same as TC neurons model previously published (14,50). The model was based on data from isolated guinea-pig hippocampal neurons, recorded at room temperature (20-22 °C) with modifications to the Boltzmann curve parameters of activation contained in the correction to the original models [59]. A small LJP (<3 mV) was not corrected (50). A *Q*_*10*_ = 3 was used for simulations at different temperatures.

### Calcium-activated potassium currents

TC neuron express genes for BK-type (54) and SK-type calcium-activated potassium channels (55). Models of BK-type currents, similar to the IC current, have already been used to model TC neurons (14,50,54). However, data characterizing this current in mammalian neurons are not available. We thus included only a model of I_SK_ (available in ModelDB, accession no. 139653) based on rat mRNA expression data in Xenopus oocytes (56).

### Intracellular calcium dynamics

A simple exponential decay mechanism was used to model the intracellular calcium dynamics (ModelDB, accession no. 139653). Both *I*_*CaT*_ and *I*_*CaL*_ contributed to the intracellular calcium concentration.

In addition, we included a voltage-insensitive membrane current *I*_*leak*_. The equilibrium potential was −79 mV and corresponded to the average resting potential from our experimental recordings.

### Simulation and parameters optimization

NEURON 7.5 software was used for simulation (57). We used NEURON variable time step method for all simulations. For the sake of spatial discretization, each section was divided into segments of 40 μm length. The following global parameters were set: initial simulation voltage (−79 mV), simulation temperature (34 °C), specific membrane capacitance (1 μF/cm^2^), specific intracellular resistivity 100 Ωcm for all the sections, equilibrium potentials for sodium and potassium were 50 mV and −90 mV, respectively.

BluePyOpt (18) with Indicator Based Evolutionary Algorithm (IBEA) were used to fit the models to the experimental data. Each optimization run was repeated with three different random seeds and evaluated 100 individuals for 100 generations. The evaluation of these 300 individuals for 100 generations was parallelized using the *iPython ipyparallel* package and took between 21 and 52 h on 48 CPU cores (Intel Xeon 2.60 GHz) on a computing cluster. Each optimization run typically resulted in tens or hundreds of unique acceptable solutions, defined as models having all feature errors below 3 STD from the experimental mean.

The models will be made publicly available at ModelDB (58). The configuration files for the optimization and analysis will be made publicly available on Github, Bluepyopt page (59).

### Sensitivity analysis

We performed a sensitivity analysis of an optimization solution by varying one parameter value (*p*_*m*_) at a time and calculating the electrical features from the voltage traces (y^+^and y^−^). We defined the sensitivity as the ratio between the normalized feature change and the parameter change, which for smooth functions approximates a partial derivative (60,61). The features changes were normalized by the optimized feature value. For small changes of parameter values, we assumed that the features depend linearly on its parameters. We could thus linearize the relationship between the features and the parameters around an optimized parameter set and calculate the derivatives. The derivatives were calculated with a central difference scheme (60).

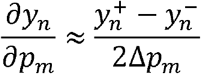

We collected the derivatives (sensitivities) in the *N* × *M* Jacobian matrix, with *N* representing the number of features and *M* the number of parameters.

To rank parameters and features we computed their relative importance by calculating their norms (the square root of the summed squared values) from the Jacobian columns and rows, respectively. To cluster parameters based on similar influences on the features and to cluster features that were similarly dependent on the parameters, we used angles between columns (or rows) to compute distances D between parameters (or features):

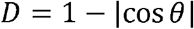

Features where thus considered similar if they depended in a similar manner on the parameters, independent of sign or magnitude.

## Acknowledgements

We thank the Blue Brain Project Visualization Team for generating the meshes and visualizing the 3D morphologies.

## Author Contributions

**Conceptualization:** E.I., S.L.H. **Data Acquisition and Curation:** J.Y., Y.S. **Formal Analysis:** E.I., J.Y., B.Z., C.O. **Funding Acquisition:** S.L.H., H.M. **Investigation:** E.I., J.Y., Y.S. **Methodology:** E.I., B.Z., C.R., W.V. **Project Administration:** C.O., S.L.H., H.M. **Resources and Software:** J.Y., Y.S., B.Z., W.V., C.R. **Supervision:** S.L.H., H.M. **Validation and Visualization:** E.I. **Writing - Original Draft Preparation:** E.I. **Writing - Review & Editing:** E.I., J.Y., C.O., W.V., H.M., S.L.H.

## Supporting Information captions

**S1 Features Dataset - Electrical features**

Spreadsheet containing experimental electrical figures, separated in three different sheets. The features, their means and standard deviations are used in Fig 1C, Fig 2, Fig 3, Fig 4C, Fig 5C, Fig 8A.

**S2 Supporting figures**

