## Supplementary Figures for "Experimentally-constrained biophysical models of tonic and burst firing modes in thalamocortical neurons" (from another version of this preprint: version 2, 2019-01-04)

### S2 Supporting figures

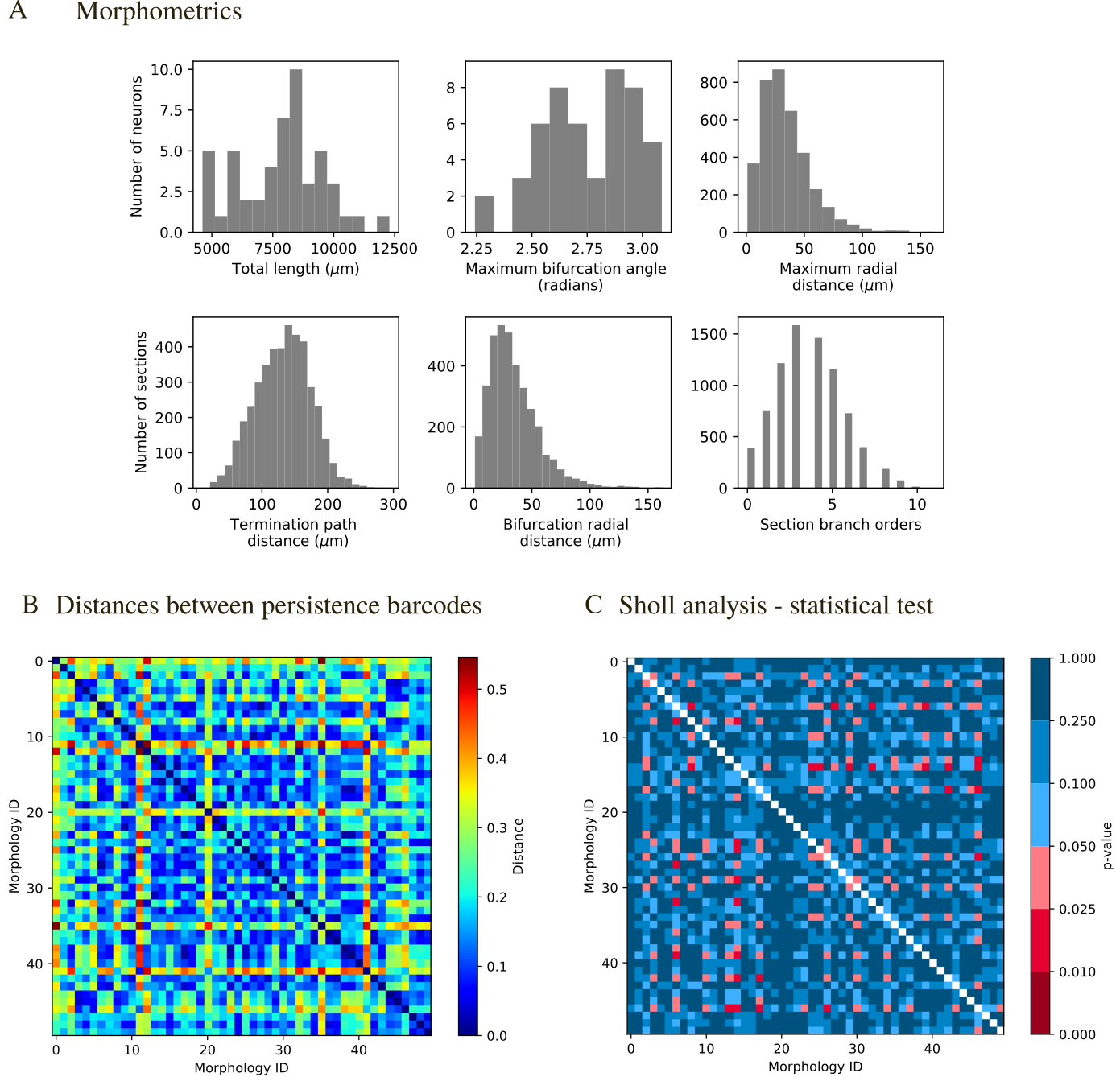

**Fig S1. Morphological analysis.**

(A) Basic morphometrics for the thalamocortical (TC) morphological type. Each histogram shows basic morphometrics at the level of the neuron (first row) or at the level of dendritic trees (second row). (B) Distance matrix between persistence barcodes for all TC morphologies. (Related to Fig 2) (C) P-values for the k-samples Anderson-Darling statistics. It tests the null hypothesis that the Sholl profiles of each pair of morphologies are drawn from the same population. The p-values are not corrected for multiple comparisons and show that we cannot reject the null hypothesis for most of the morphology pairs (at 0.05 significance level).

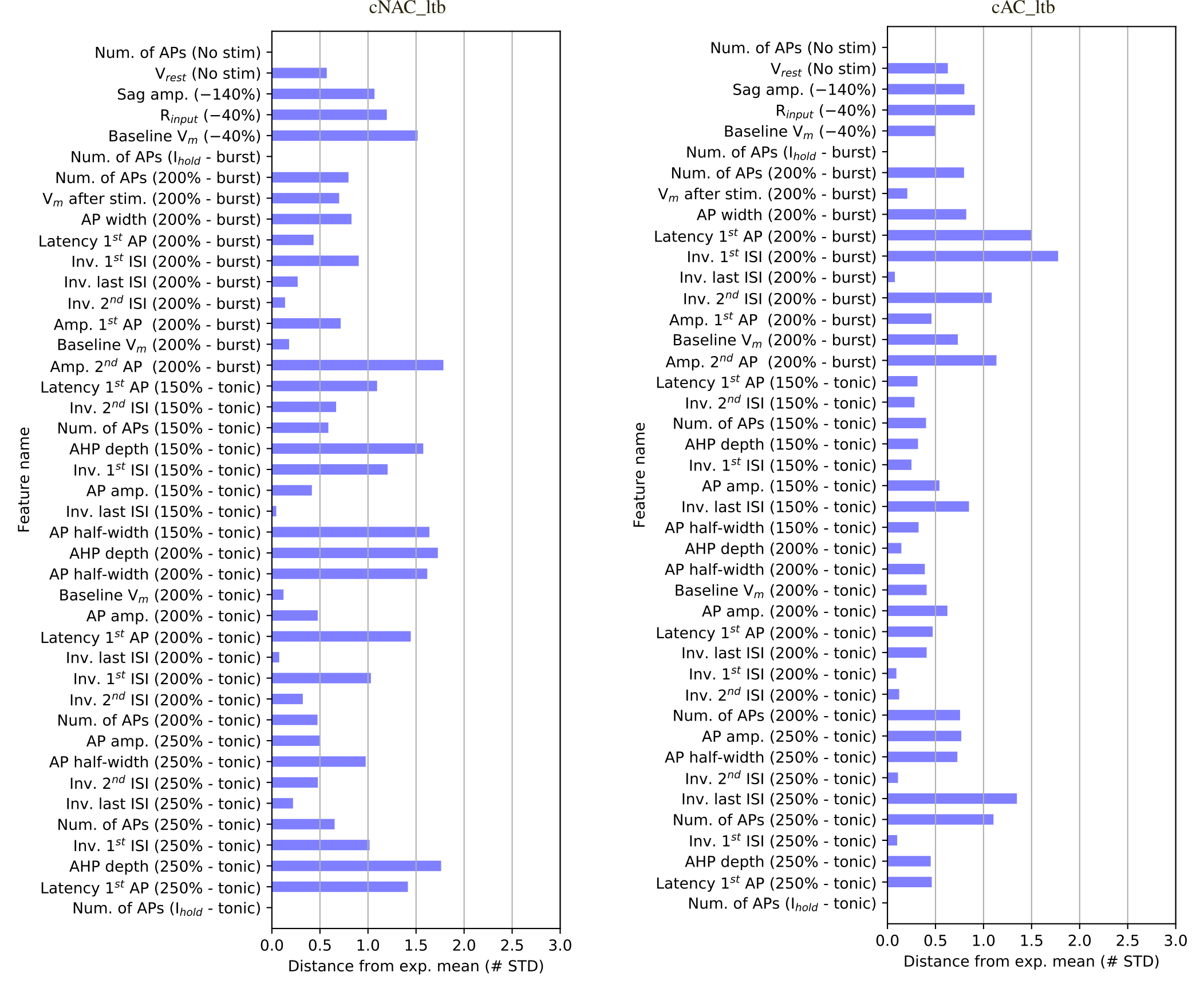

**Fig S2. Feature errors for the models shown in Fig 4**. Where the error bar is missing the error value is 0.

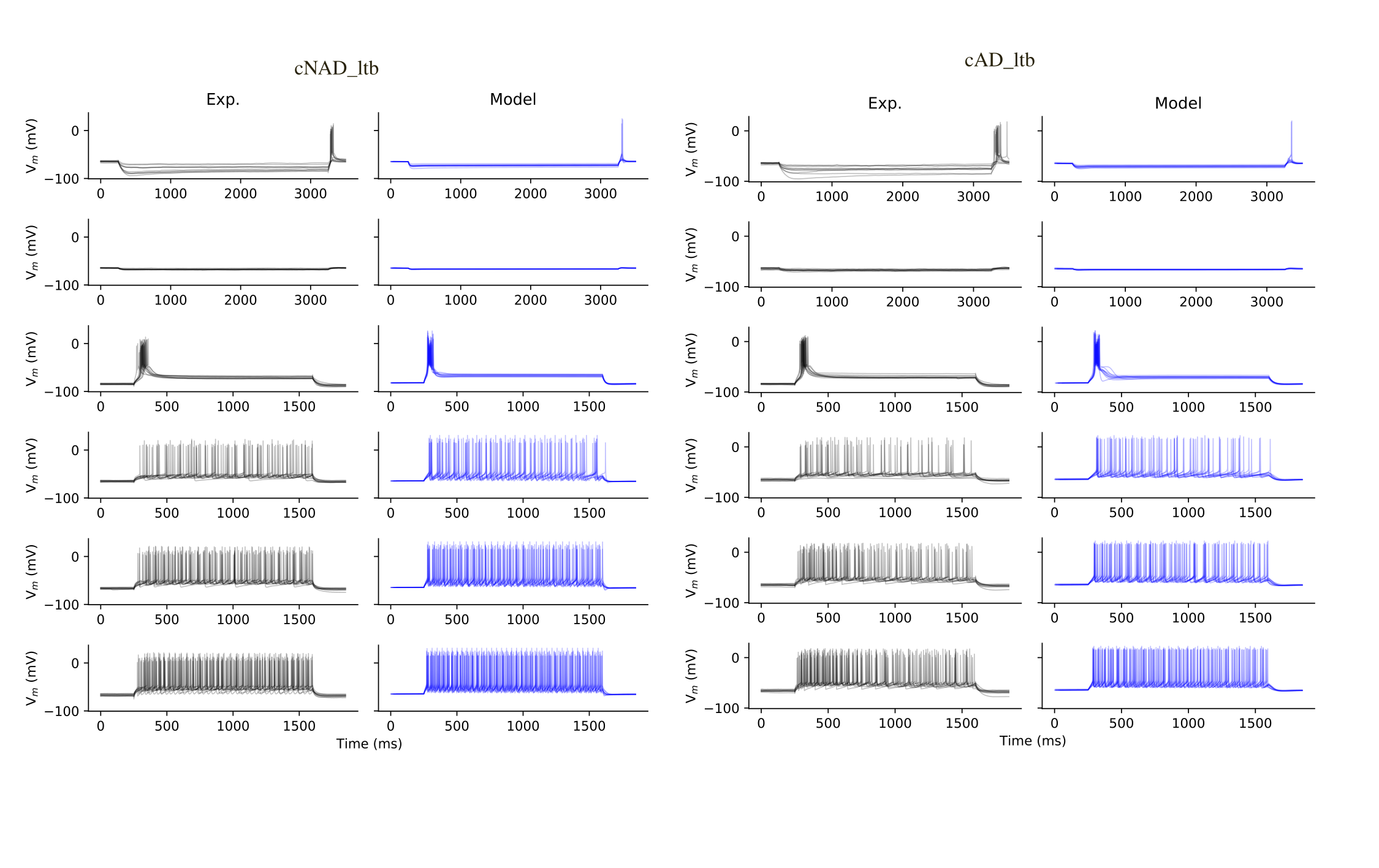

**Fig S3. Example of traces from experimental recordings and models.** Voltage traces from a sample of 10 different experimental cells and models, corresponding to the features shown in Fig 5.

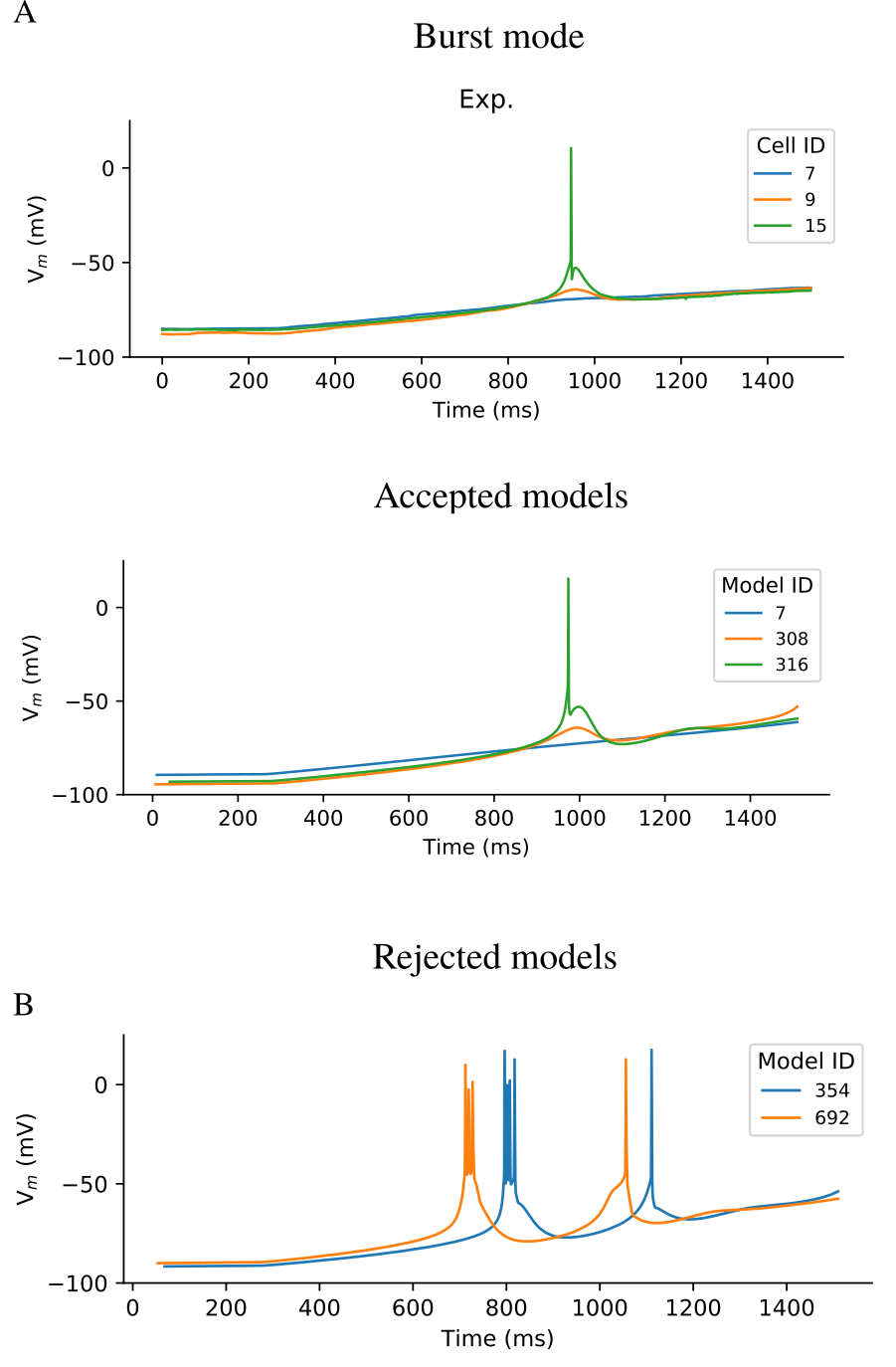

**Fig S4. Variability of experimental and model voltage responses to ramp stimuli.** (A) Experimental cells and models show four different types of behavior, when recorded in burst mode: no firing, only low threshold spike, low-threshold burst (low-threshold spikes crowned by sodium spikes), low-threshold burst followed by tonic firing.
